# Quasi-essentiality of RNase Y in *Bacillus subtilis* is caused by its critical role in the control of mRNA homeostasis

**DOI:** 10.1101/2020.05.20.106237

**Authors:** Martin Benda, Simon Woelfel, Katrin Gunka, Stefan Klumpp, Anja Poehlein, Debora Kálalová, Hana Šanderová, Rolf Daniel, Libor Krásný, Jörg Stülke

## Abstract

RNA turnover is essential in all domains of life. The endonuclease RNase Y (*rny*) is one of the key components involved in RNA metabolism of the model organism *Bacillus subtilis*. Essentiality of RNase Y has been a matter of discussion, since deletion of the *rny* gene is possible, but leads to severe phenotypic effects. In this work, we demonstrate that the *rny* mutant strain rapidly evolves suppressor mutations to at least partially alleviate these defects. All suppressor mutants had acquired a duplication of an about 60 kb long genomic region encompassing genes for all three core subunits of the RNA polymerase – α, β, β′. When the duplication of the RNA polymerase genes was prevented by relocation of the *rpoA* gene in the *B. subtilis* genome, all suppressor mutants carried distinct single point mutations in evolutionary conserved regions of genes coding either for the β or β’ subunits of the RNA polymerase that were not tolerated by wild type bacteria. *In vitro* transcription assays with the mutated polymerase variants showed massive decreases in transcription efficiency. Altogether, our results suggest a tight cooperation between RNase Y and the RNA polymerase to establish an optimal RNA homeostasis in *B. subtilis* cells.

## INTRODUCTION

Among all organisms, bacteria are the ones multiplying most rapidly. Under optimal conditions, the model bacteria *Escherichia coli* and *Bacillus subtilis* have generation times of 20 to 30 minutes. On the other hand, bacteria are exposed to a variety of changing environmental conditions, and due to their small size, the impact of environmental changes is particularly severe for bacterial cells. To adapt to these potentially rapidly changing conditions, bacteria have evolved a huge arsenal of systems to sense and respond to the environment. Especially in the competition between microorganisms, it is crucial that these responses are both rapid and productive. However, while regulatory events may be very rapid, there is an element of retardation in the system, and this is the stability of mRNA and protein molecules. If the continued activity of a protein may become harmful to the bacteria, it is important not only to prevent expression of the corresponding gene but also to take two important measures: (i) switch off the protein’s activity and (ii) degrade the mRNA to exclude further production of the protein. The inactivation or even degradation of proteins is well documented in the model bacteria. For example, in both *E. coli* and *B. subtilis* the uptake of toxic ammonium is limited by a regulatory interaction of the ammonium transporter with GlnK, a regulatory protein of the PII family (1,2). Similarly, the uptake of potentially toxic potassium can be prevented by inhibition of potassium transporters at high environmental potassium concentrations, either by the second messenger cyclic di-AMP or by interaction with a dedicated modified signal transduction protein, PtsN (3,4,5). To prevent the accumulation of potentially harmful mRNAs, bacteria rely on a very fast mRNA turnover. Indeed, in *E. coli* and *B. subtilis* more than 80% of all transcripts have average half-lives of less than 8 minutes, as compared to about 30 minutes and 10 hours in yeast or human cells, respectively (6,7,8,9). Thus, the mRNA turnover is much faster than the generation time. The high mRNA turnover rate in bacteria contributes to the fast adaptation even in rapidly growing cells. The rapid mRNA turnover is therefore a major factor to resolve the apparent growth speed-adaptation trade-off.

RNases are the key elements to achieve the rapid mRNA turnover in bacteria. Theses enzymes can degrade bulk mRNA in a rather unspecific manner, just depending on the accessibility of the RNA molecules as well as perform highly specific cleavages that serve to process an RNA molecule to its mature form. In all organisms, RNA degradation involves an interplay of endo- and exoribonucleases as well as other proteins such as RNA helicases that resolve secondary structures (10,11,12,13). Often, these proteins form a complex called the RNA degradosome. In *E. coli*, the RNA degradosome is organized around the essential endoribonuclease RNase E (14,15). RNase E consists of two parts, the N-terminal endoribonuclease domain that harbors the enzymatic activity and the C-terminal macromolecular interaction domain that serves as the scaffold for the degradosome components and is responsible for the binding of RNase E to the cell membrane (15,16). As mentioned above, RNase E is essential for viability of the bacteria. An analysis of the contributions of the two parts of RNase E to its essentiality revealed that the enzymatically active N-terminal domain is essential whereas the C-terminal interaction domain is dispensable (17). This suggests that the endoribonucleolytic attack on mRNA molecules is the essential function of RNase E, whereas the interaction with other degradosome components is not required for viability. This conclusion is supported by the fact, that the other components of the *E. coli* degradosome are also dispensable (14).

RNase E is widespread in proteobacteria, cyanobacteria, and actinobacteria, but absent from many firmicutes, ε-proteobacteria, or from bacteria of the *Deinococcus*-*Thermus* class. However, an efficient RNA-degrading machinery is important also for these bacteria to allow both rapid growth and adaptation. Indeed, these bacteria possess a different endoribonuclease, RNase Y (18,19). A depletion of RNase Y results in a two-fold increase of the average mRNA half-life in *B. subtilis* (19). Similar to RNase E, RNase Y is a membrane protein, and it is capable of interacting with several proteins involved in RNA degradation. Among these proteins are the 5′-to-3′ exoribonunclease RNase J1, polynucleotide phosphorylase, the RNA helicase CshA, the glycolytic proteins enolase and phosphofructokinase, and a protein complex composed of YaaT, YlbF, and YmcA (18,19,20,21,22,23). Many of these interactions are likely to be transient as judged from the distinct localization of RNase Y and its interaction partners in the cell membrane and in the cytoplasm, respectively (24).

We are interested in the identification of the essential cellular components that are required for the viability of *B. subtilis* cells with the aim to construct strains that harbor only the minimal set of genes to fulfill the essential cellular functions (25,26,27). For *B. subtilis*, RNase Y and RNase J1 were originally described as being essential (18,19,28,29,30). Interestingly, these two RNases are also present in the most genome-reduced independently viable organism, *Mycoplasma mycoides* JCVI-syn3.0 (31). Both RNase J1 and RNase Y are involved in the processing and degradation of a large number of RNA molecules in *B. subtilis* (32,33,34,35,36). However, more recent studies demonstrated the possibility to delete the *rnjA* and *rny* genes, encoding the two RNases (37,38). The dispensability of RNase Y was confirmed in a global approach to inactivate all genes of *B. subtilis*. However, an *rnjA* mutant could not be obtained in this latter study (39).

Comprehensive knowledge on essential genes and functions is the key to construct viable minimal genomes. By definition, essential genes cannot be individually deleted in a wild type genetic background under standard growth conditions (25). In this study, we have addressed the essentiality of RNase Y in *B. subtilis*. While the *rny* gene could indeed be deleted, this was accompanied by the rapid acquisition of suppressor mutations that affect the transcription apparatus. We demonstrate that a strongly reduced transcription activity is required to allow stable growth of *B. subtilis* in the absence of RNase Y. Our results suggest that the accumulation of mRNA that cannot be degraded is the growth-limiting factor in strains lacking RNase Y.

## MATERIAL AND METHODS

### Bacterial strains, plasmids and growth conditions

All *B. subtilis* strains used in this study are listed in Table 1. All strains are derived from the laboratory strain 168 (*trpC2*). *B. subtilis* and *E. coli* cells were grown in Lysogeny Broth (LB medium) (40). LB plates were prepared by addition of 17 g Bacto agar/l (Difco) to LB (40). The plasmids are listed in Table 2. Oligonucleotides are listed in Table S1.

**Table 1.**
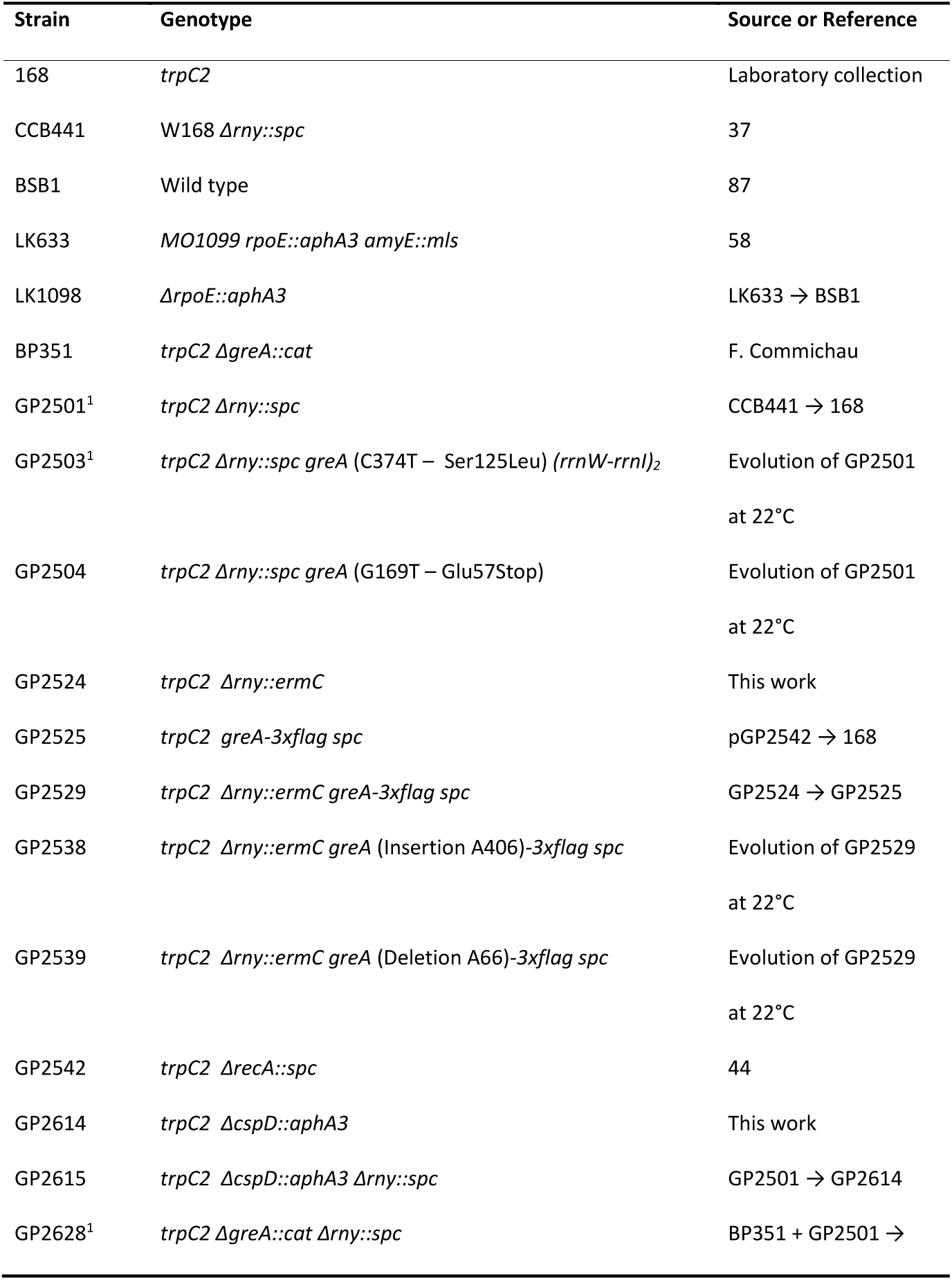
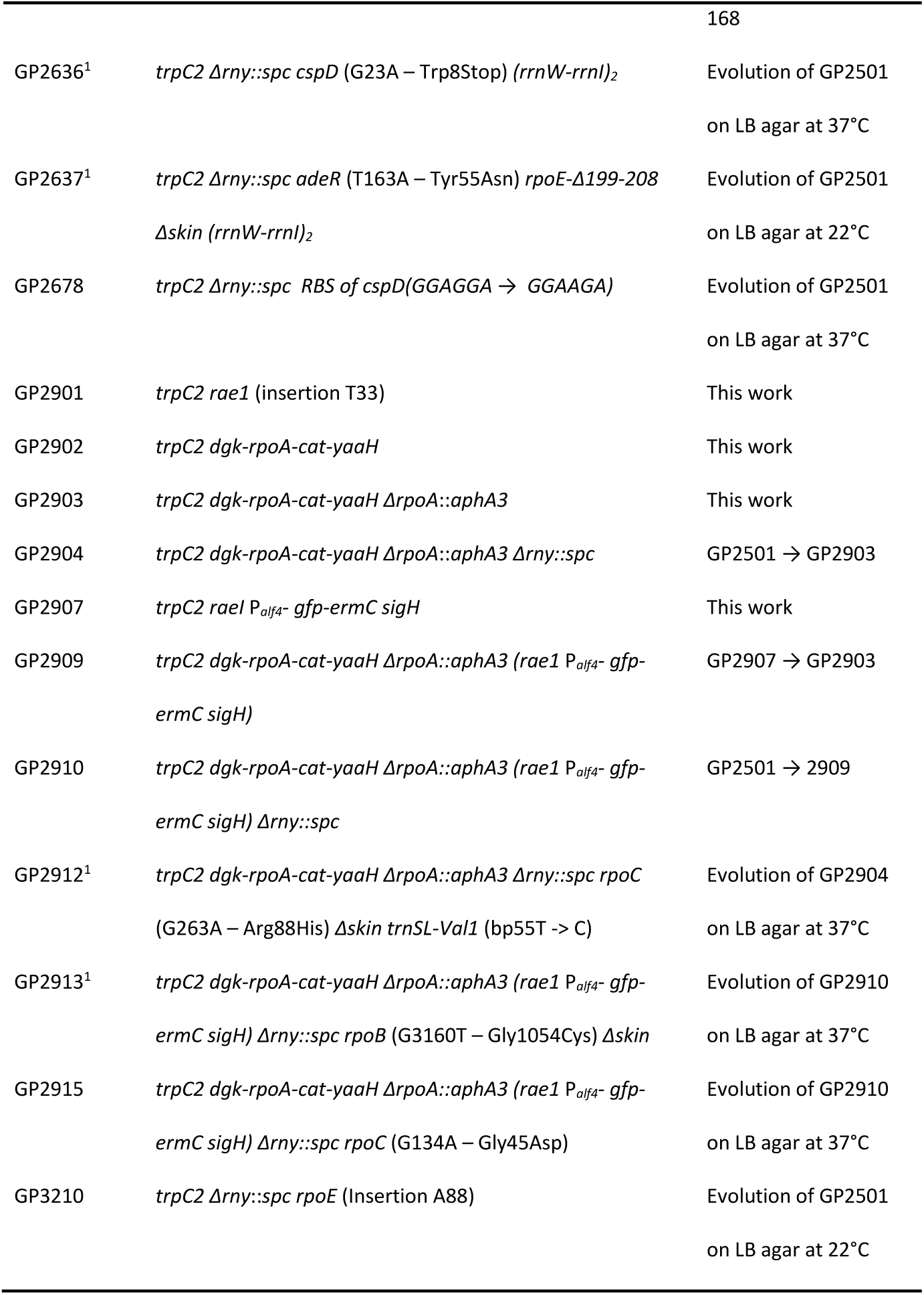
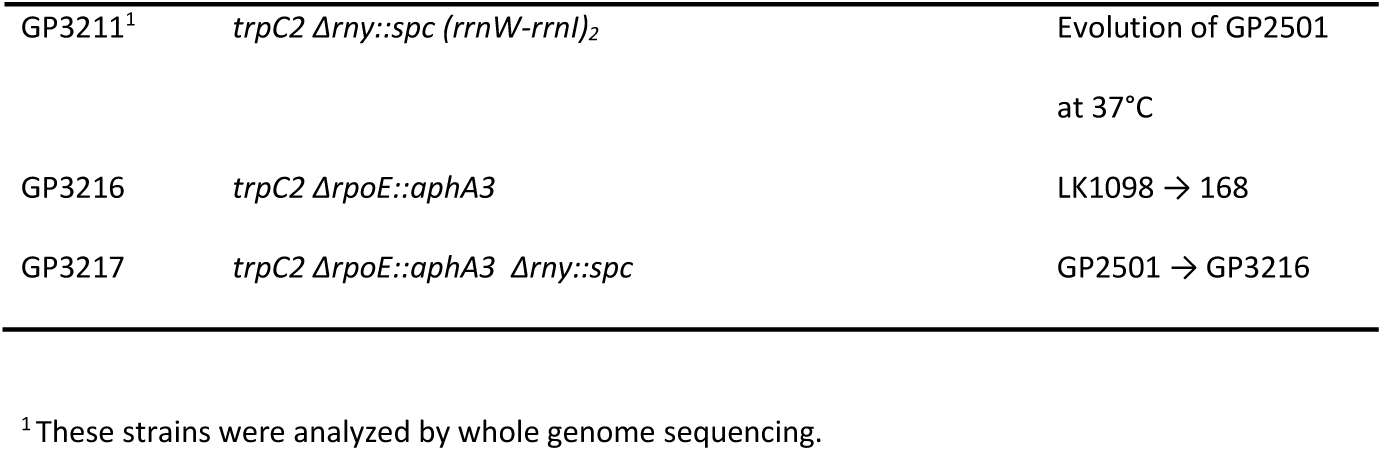
*B. subtilis* strains used in this study

| Strain | Genotype | Source or Reference |
| --- | --- | --- |
| 168 | *trpC2* | Laboratory collection |
| CCB441 | W168 $\Delta rny::spc$ | 37 |
| BSB1 | Wild type | 87 |
| LK633 | MO1099 *rpoE::aphA3 amyE::mls* | 58 |
| LK1098 | $\Delta rpoE::aphA3$ | LK633 → BSB1 |
| BP351 | *trpC2* $\Delta greA::cat$ | F. Commichau |
| GP2501<sup>1</sup> | *trpC2* $\Delta rny::spc$ | CCB441 → 168 |
| GP2503<sup>1</sup> | *trpC2* $\Delta rny::spc greA$ (C374T – Ser125Leu) (*rrnW-rrnI*)<sub>2</sub> | Evolution of GP2501<br>at 22°C |
| GP2504 | *trpC2* $\Delta rny::spc greA$ (G169T – Glu57Stop) | Evolution of GP2501<br>at 22°C |
| GP2524 | *trpC2* $\Delta rny::ermC$ | This work |
| GP2525 | *trpC2 greA-3xflag spc* | pGP2542 → 168 |
| GP2529 | *trpC2* $\Delta rny::ermC greA-3xflag spc$ | GP2524 → GP2525 |
| GP2538 | *trpC2* $\Delta rny::ermC greA$ (Insertion A406)-3xflag *spc* | Evolution of GP2529<br>at 22°C |
| GP2539 | *trpC2* $\Delta rny::ermC greA$ (Deletion A66)-3xflag *spc* | Evolution of GP2529<br>at 22°C |
| GP2542 | *trpC2* $\Delta recA::spc$ | 44 |
| GP2614 | *trpC2* $\Delta cspD::aphA3$ | This work |
| GP2615 | *trpC2* $\Delta cspD::aphA3 \Delta rny::spc$ | GP2501 → GP2614 |
| GP2628<sup>1</sup> | *trpC2* $\Delta greA::cat \Delta rny::spc$ | BP351 + GP2501 → |
|  |  | 168 |
| GP2636<sup>1</sup> | *trpC2 Δrny::spc cspD* (G23A – Trp8Stop) (*rrnW-rrnI*)<sub>2</sub> | Evolution of GP2501<br>on LB agar at 37°C |
| GP2637<sup>1</sup> | *trpC2 Δrny::spc adeR* (T163A – Tyr55Asn) *rpoE-Δ199-208*<br>*Δskin (rrnW-rrnI)*<sub>2</sub> | Evolution of GP2501<br>on LB agar at 22°C |
| GP2678 | *trpC2 Δrny::spc* RBS of *cspD*(GGAGGA → GGAAGA) | Evolution of GP2501<br>on LB agar at 37°C |
| GP2901 | *trpC2 rae1* (insertion T33) | This work |
| GP2902 | *trpC2 dgk-rpoA-cat-yaaH* | This work |
| GP2903 | *trpC2 dgk-rpoA-cat-yaaH ΔrpoA::aphA3* | This work |
| GP2904 | *trpC2 dgk-rpoA-cat-yaaH ΔrpoA::aphA3 Δrny::spc* | GP2501 → GP2903 |
| GP2907 | *trpC2 rae1 P<sub>alf4</sub>- gfp-ermC sigH* | This work |
| GP2909 | *trpC2 dgk-rpoA-cat-yaaH ΔrpoA::aphA3 (rae1 P<sub>alf4</sub>- gfp-ermC sigH)* | GP2907 → GP2903 |
| GP2910 | *trpC2 dgk-rpoA-cat-yaaH ΔrpoA::aphA3 (rae1 P<sub>alf4</sub>- gfp-ermC sigH) Δrny::spc* | GP2501 → 2909 |
| GP2912<sup>1</sup> | *trpC2 dgk-rpoA-cat-yaaH ΔrpoA::aphA3 Δrny::spc rpoC*<br>(G263A – Arg88His) *Δskin trnSL-Val1* (bp55T → C) | Evolution of GP2904<br>on LB agar at 37°C |
| GP2913<sup>1</sup> | *trpC2 dgk-rpoA-cat-yaaH ΔrpoA::aphA3 (rae1 P<sub>alf4</sub>- gfp-ermC sigH) Δrny::spc rpoB* (G3160T – Gly1054Cys) *Δskin* | Evolution of GP2910<br>on LB agar at 37°C |
| GP2915 | *trpC2 dgk-rpoA-cat-yaaH ΔrpoA::aphA3 (rae1 P<sub>alf4</sub>- gfp-ermC sigH) Δrny::spc rpoC* (G134A – Gly45Asp) | Evolution of GP2910<br>on LB agar at 37°C |
| GP3210 | *trpC2 Δrny::spc rpoE* (Insertion A88) | Evolution of GP2501<br>on LB agar at 22°C |
| GP3211<sup>1</sup> | *trpC2 Δrny::spc (rrnW-rrnI)<sub>2</sub>* | Evolution of GP2501<br>at 37°C |
| GP3216 | *trpC2 ΔrpoE::aphA3* | LK1098 → 168 |
| GP3217 | *trpC2 ΔrpoE::aphA3 Δrny::spc* | GP2501 → GP3216 |

<sup>1</sup>These strains were analyzed by whole genome sequencing.

**Table 2.** Plasmids used in this study

| Plasmid | Relevant characteristics | Source or Reference |
| --- | --- | --- |
| pCD2 | For overexpression of *B. subtilis* $\sigma^A$ | 54 |
| pJOE8999 | CRISPR-Cas9 vector | 49 |
| pBSURNAP | *P<sub>T7</sub> rpoA rpoZ rpoE rpoY rpoB-rpoC-8xHis* | This work |
| pGP1331 | Allows construction of triple FLAG-tag fusions | 88 |
| pGP2181 | *P<sub>T7</sub> rpoA rpoZ rpoE rpoY rpoB-rpoC\*-8xHis* (RpoC-R88H) | This work |
| pGP2182 | *P<sub>T7</sub> rpoA rpoZ rpoE rpoY rpoB\*-rpoC-8xHis* (RpoB-G1054C) | This work |
| pGP2542 | pGP1331/ *greA-3xflag spc* | This work |
| pGP2825 | pJOE8999/ *rpoC* (G263A) | This work |
| pGP2826 | pJOE8999/*rea1* (insertion T33) | This work |
| pRLG770 | promoter vector | 89 |
| pRLG7558 | pRLG770 with *B. subtilis* *P<sub>veg</sub>* (-38/-1, +1G) | 68 |
| pRLG7596 | pRLG770 with *B. subtilis* *rrnB* P1 (-39/+1) | 68 |
| pLK502 | pRLG770 with *B. subtilis* *P<sub>ilvB</sub>* (-262/-1, +1GG) | This work |

### DNA manipulation and genome sequencing

*B. subtilis* was transformed with plasmids, genomic DNA or PCR products according to the two-step protocol (40,41). Transformants were selected on LB plates containing erythromycin (2 µg/ml) plus lincomycin (25 µg/ml), chloramphenicol (5 µg/ml), kanamycin (10 µg/ml), or spectinomycin (250 µg/ml). Competent cells of *E. coli* were prepared and transformed following the standard procedure (40) and selected on LB plates containing kanamycin (50 µg/ml). S7 Fusion DNA polymerase (Mobidiag, Espoo, Finland) was used as recommended by the manufacturer. DNA fragments were purified using the QIAquick PCR Purification Kit (Qiagen, Hilden, Germany). DNA sequences were determined by the dideoxy chain termination method (40). Chromosomal DNA from *B. subtilis* was isolated using the peqGOLD Bacterial DNA Kit (Peqlab, Erlangen, Germany). To identify the mutations in the suppressor mutant strains GP2503, GP2636, GP2637, GP2912, GP2913, and GP3211 (see Table 1), the genomic DNA was subjected to whole-genome sequencing. Concentration and purity of the isolated DNA was first checked with a Nanodrop ND-1000 (PeqLab Erlangen, Germany) and the precise concentration was determined using the Qubit® dsDNA HS Assay Kit as recommended by the manufacturer (Life Technologies GmbH, Darmstadt, Germany). Illumina shotgun libraries were prepared using the Nextera XT DNA Sample Preparation Kit and subsequently sequenced on a MiSeq system with the reagent kit v3 with 600 cycles (Illumina, San Diego, CA, USA) as recommended by the manufacturer. The reads were mapped on the reference genome of *B. subtilis* 168 (GenBank accession number: NC_000964) (42).

Mapping of the reads was performed using the Geneious software package (Biomatters Ltd., New Zealand) (43). Frequently occurring hitchhiker mutations (44) and silent mutations were omitted from the screen. The resulting genome sequences were compared to that of our in-house wild type strain. Single nucleotide polymorphisms were considered as significant when the total coverage depth exceeded 25 reads with a variant frequency of ≥90%. All identified mutations were verified by PCR amplification and Sanger sequencing. Copy numbers of amplified genomic regions were determined by dividing the mean coverage of the amplified regions by the mean coverage of the remaining genome (45).

### Construction of deletion mutants

Deletion of the *rny, rpoA*, and *cspD* genes was achieved by transformation with PCR products constructed using oligonucleotides to amplify DNA fragments flanking the target genes and intervening antibiotic resistance cassettes as described previously (46,47,48). The identity of the modified genomic regions was verified by DNA sequencing.

### Chromosomal relocation of the *rpoA* gene

To construct a strain in which the genes for the core subunits of RNA polymerase are genomically separated, we decided to place the *rpoA* gene between the *dgk* and *yaaH* genes, and then to delete the original copy of the gene. First, the *rpoA* gene was fused in a PCR reaction with its cognate promoter and a chloramphenicol resistance gene at the 5**′** and 3**′** ends, respectively. In addition, the amplified *dgk* and *yaaH* genes were fused to this construct to direct the integration of the construct to the *dgk-yaaH* locus. The fusion of PCR products was achieved by overlapping primers. The final product was then used to transform *B. subtilis* 168. Correct insertion was verified by PCR amplification and sequencing. The resulting strain was *B. subtilis* GP2902. In the second step, the original *rpoA* gene was replaced by a kanamycin resistance gene as described above, leading to strain GP2903.

### Genome editing

Introduction of genetic changes in genes for RNA polymerase subunit RpoC or the non-essential RNase Rae1 at their native locus was attempted using CRISPR editing as described (49). Briefly, oligonucleotides encoding a 20 nucleotide gRNA with flanking *BsaI* sites and a repair fragment carrying mutations of interest with flanking *SfiI* restriction sites were cloned sequentially into vector pJOE8999 (49). The resulting plasmids pGP2825 and pGP2826 were used to transform recipient *B. subtilis* strain 168 and cells were plated on 10 μg/ml kanamycin plates with 0.2% mannose. Transformation was carried out at 30°C since replication of pJOE8999 derivatives is temperature-sensitive. The transformants were patched on LB agar plates and incubated at the non-permissive temperature of 50°C. The loss of the vector was verified by the inability of the bacteria to grow on kanamycin plates. The presence of the desired mutation in *rae1* or *rpoC* was checked via Sanger sequencing. While the desired mutation could be introduced into the *rae1* gene, this was not the case for *rpoC*.

### Construction of the expression vector pBSURNAP

To facilitate the purification of different variants of *B. subtilis* RNA polymerase, we expressed and purified the core subunits of the RNA polymerase and the sigma factor separately in *E. coli*. For the expression of the core subunits, we cloned the corresponding *B. subtilis* genes into the backbone of a pET28a derivative as follows. The pRMS4 vector (a pET28a derivative, 50) containing *Mycobacterium smegmatis* RNA polymerase core subunit genes was used as a template to create an analogous vector containing the genes *rpoA, rpoZ, rpoE, rpoY*, and *rpoBC*. The construct was designed to allow removal/substitution of each gene via unique restriction sites (Fig. S1). DNA encoding *rpo*A, *rpo*Z, *rpo*E and *rpo*Y genes was cloned as one single fragment (purchased as Gene Art Strings from Invitrogen) via XbaI and NotI restriction sites. The *rpoB* and *rpoC* genes were amplified by PCR using genomic DNA of *B. subtilis* 168 as a template and inserted into the plasmid via NotI and NcoI or NcoI and KpnI restriction sites, respectively. The rpoC gene was inserted with a sequence encoding a 8xHis tag on the 3’ end. The cloned construct was verified by DNA sequencing. The final vector, pBSURNAP, encodes a polycistronic transcript for expression of all six RNA polymerase core subunits. Expression is driven from an IPTG-inducible T7 RNAP-dependent promoter. Each gene is preceded by a Shine-Dalgarno sequence (AGGAG) except for *rpoC*. RpoB-RpoC are expressed as one fused protein connected by a short linker (9 amino acid residues) to decrease the possibility that *E. coli* subunits would mix with *B. subtilis* subunits as done previously for RNA polymerase from *Mycobacterium bovis* (51). The full sequence of pBSURNAP has been deposited in GenBank under Accession No. MT459825. The mutant alleles of *rpoB* and *rpoC* were amplified from the mutant strains GP2913 and GP2912 and introduced into pBSURNAP by replacing the wild type alleles as NotI/NcoI and NcoI/KpnI fragments, respectively. The resulting plasmids were pGP2181 (RpoC-R88H) and pGP2182 (RpoB-G1054C).

### Purification of *B. subtilis* RNA polymerase from *E. coli* cells

For purification, *E. coli* BL21 carrying pBSURNAP or the plasmids specifying the mutant alleles was cultivated in LB medium containing kanamycin (50 µg/ml). Expression was induced by the addition of IPTG (final concentration 0.3 mM) to logarithmically growing cultures (OD600 between 0.6 and 0.8), and cultivation was continued for three hours. Cells were harvested and the pellets from 1 l of culture medium were washed in 50 ml buffer P (300 mM NaCl, 50 mM Na2HPO4, 3 mM β-mercaptoethanol, 1 mM PMSF, 5% glycerol) and the pellets were resuspended in 30 ml of the same buffer. Cells were lysed using a HTU DIGI-F Press (18,000 p.s.i., 138,000 kPa, two passes, G. Heinemann, Germany). After lysis, the crude extracts were centrifuged at 41,000 x *g* for 30 min at 4°C, and the RNA polymerase was purified from the supernatant via the His-tagged RpoC as described (52). The RNA polymerase-containing fractions were pooled and further purified by size exclusion chromatography. For this purpose, the complex was applied onto a HiLoad 16/600 Superdex 200 column (GE Healthcare) in buffer P. The buffer was filtered (0.2 µm filters) prior to protein separation on an Äkta Purifier (GE Healthcare). The fractions containing RNA polymerase were pooled and dialyzed against RNA polymerase storage buffer (50 mM Tris–HCl, pH 8.0, 3 mM β-mercaptoethanol, 0.15 M NaCl, 50% glycerol, 1:1,000). The purified RNA polymerase was stored at −20°C.

The housekeeping sigma factor σ^A^ was overproduced from plasmid pCD2 (53) and purified as described (54).

### *In vitro* transcription assays

Multiple round transcription assays were performed as described previously (55), unless stated otherwise. Initiation competent RNA polymerase was reconstituted using the core enzyme and saturating concentration of σ^A^ in dilution buffer (50 mM Tris-HCl, pH 8.0, 0.1 M NaCl, 50% glycerol) for 10 min at 30°C. Assays were carried out in 10 μl with 64 nM RNA polymerase holoenzyme and 100 ng plasmid DNA templates in transcription buffer containing 40 mM Tris-HCl (pH 8.0), 10 mM MgCl2, 1 mM dithiothreitol (DTT), 0.1 mg/ml bovine serum albumin (BSA), 150 mM NaCl, and NTPs (200 μM ATP, 2,000 μM GTP, 200 μM CTP, 10 μM UTP plus 2 μM of radiolabeled [α-^32^P]-UTP). The samples were preheated for 10 min at 37°C. The reaction was started by the addition of RNA polymerase and allowed to proceed for 20 min (30 min in the case of iNTP-sensing experiments) at 37°C. Subsequently, the reaction was stopped by the addition of 10 μl of formamide stop solution (95% formamide, 20 mM EDTA, pH 8.0). The samples were loaded onto 7M urea-7% polyacrylamide gels. The gels were dried and exposed to Fuji MS phosphor storage screens, scanned with a Molecular Imager FX (BIORAD) and analyzed with Quantity One program (BIORAD).

## RESULTS

### Inactivation of the *rny* gene leads to fast evolution of suppressor mutations affecting transcription

RNase Y had been considered to be essential (18,28); however, two studies reported that the *rny* gene could be deleted from the genome (37,39). The deletion leads to severe growth defects and morphological changes (37). In an attempt to get a better understanding of the importance of RNase Y for *B. subtilis* physiology, we deleted the *rny* gene in the genetic background of *B. subtilis* 168. The colonies of the resulting strain, GP2501, were small and lysed rapidly. Moreover, the cells grew very slowly at low temperatures (below 22°C). However, we observed the rapid appearance of suppressor mutants. By analysis of such mutants we wished to gain a better understanding of the growth-limiting problem of the *rny* mutant. For this purpose, we isolated suppressor mutants in two different experimental setups. First, suppressor mutants were adapted to growth in liquid LB medium at 22°C since the *rny* mutants had a severe growth defect at low temperatures. Indeed, the *rny* mutant GP2501 was essentially unable to grow at 22°C. After the adaptation experiment, the culture was plated at 22°C, and two colonies were isolated for further investigation. Growth of the isolated strains was verified (Fig. 1A), and one of them (GP2503) was subjected to whole genome sequencing. This confirmed the deletion of the *rny* gene and revealed the presence of an additional mutation that resulted in an amino acid substitution (S125L) in the *greA* gene encoding a transcription elongation factor (56). For the other suppressor mutant (GP2504), we sequenced the *greA* gene to test whether it had also acquired a mutation in this gene. Indeed, we found a different mutation in *greA*, resulting in the introduction of a premature stop codon after E56. Moreover, we evolved two additional suppressor mutants applying this adaptive scenario, and both contained frameshift mutations in *greA* that resulted in premature stop codons after amino acid 23 and 137 (GP2539 and GP2538, respectively; see Table 1). In conclusion, the inactivation of the *greA* gene seems to be the major suppressing mechanism using this adaptive selection scheme.

**Figure 1.**
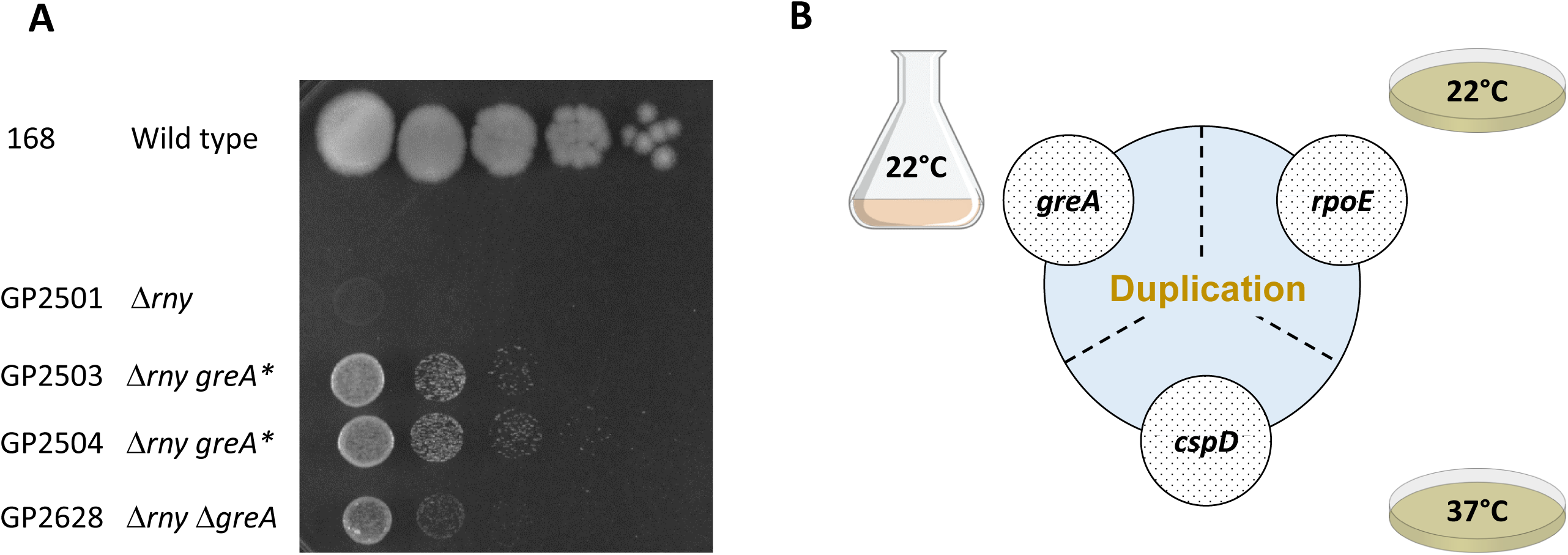
Suppressors of *rny* show increased growth at 22°C. (A) Serial drop dilutions comparing growth of the wild type strain 168, the *rny* mutant GP2501, its *greA* suppressors (GP2503, GP2504, see Table 1 for the precise mutations) and the *rny greA* double mutant GP2628 on LB-agar plate at 22°C. The picture was taken after 2 days of incubation. (B) Schematic depiction of different single nucleotide polymorphisms identified in the initial suppressor screen and their overlap with the duplication of *ctsR-pdaB* region.

Second, in addition to the adaptation experiment in liquid medium, we also evolved suppressors on solid-LB agar plates both at 22°C and 37°C. We isolated two mutants under each condition. For each temperature, one mutant was analysed by whole genome sequencing. The strain isolated at 22°C (GP2637) had a deletion of the skin element, an amino acid substitution (Y55N) in the AdeR activator protein (57), and a short internal deletion in the *rpoE* gene encoding the δ subunit of RNA polymerase, which resulted in a frameshift after residue G66 (54,58). For the second mutant isolated at 22°C (GP3210), we re-sequenced the *adeR* and *rpoE* genes. While the *adeR* gene was identical to the wild type, we found an insertion of an adenine residue after position 87 of *rpoE*, resulting in a frameshift after 29 amino acids and premature stop codon after 38 amino acids. The suppressor evolved at 37°C (GP2636) contained a mutation resulting in the introduction of a premature stop codon at position 8 in the *cspD* gene encoding an RNA binding protein which has antitermination activity in *E. coli* (59,60). Sanger sequencing of the second suppressor isolated under the same condition (GP2678) also identified a mutation affecting *cspD*, but this time in its ribosomal binding site (GGAGGA → GGAAGA). Thus, the selective pressure on agar plates at 22°C and 37°C was directed at the inactivation of the RNA polymerase subunit RpoE or the RNA binding protein CspD, respectively.

Taken together, we found mutations affecting transcription in every single suppressor mutant analysed, i. e. *greA, rpoE*, and *cspD*. It is therefore tempting to speculate that the inactivation of these genes is causative for the suppression.

### Suppression of the loss of RNase Y requires the duplication of a chromosomal region encompassing the genes encoding RNA polymerase

In order to test whether the inactivation of the *greA, rpoE*, or *cspD* genes is sufficient for the suppression of the *rny* mutant strain, we constructed the corresponding double mutants. As both *rny* and *greA* mutants are defective in genetic competence (39), the *greA rny* double mutant was obtained by transforming the wild type strain 168 with DNA molecules specifying both deletions simultaneously (see Table 1). In contrast to our expectations, the double mutants did not phenocopy the original suppressor mutants, instead the gene deletions conferred only partial suppression (for the *rny greA* double mutant, see Fig. 1A) or even no suppression (for the *rny cspD* and *rny rpoE* double mutants, data not shown). Thus, we assumed that the suppressor strains might carry additional mutations that had escaped our attention.

Indeed, a re-evaluation of the genome sequences revealed that in addition to the distinct point mutations described above there was one feature common for all the suppressors tested, regardless of the isolation condition, which was not present in the progenitor strain GP2501: It was a genomic duplication of the approximately 60 kb long *ctsR-pdaB* region. This genomic segment is flanked by clusters of ribosomal RNA operons. Upstream of the duplicated region are the *rrnJ* and *rrnW* operons, and downstream the *rrnI, rrnH*, and *rrnG* operons. This duplicated region contains 76 genes encoding proteins of various functions, among them proteolysis (ClpC), signal transduction (DisA), RNA modification (YacO, TruA), RNases (MrnC, Rae1), translation factors (EF-G, IF-1, EF-Tu), several ribosomal proteins, and proteins involved in transcription (NusG, RpoA, RpoB, RpoC, SigH). Strikingly, the genes for all three main subunits of the RNA polymerase – *rpoA, rpoB* and *rpoC* were included in the duplicated region. MrnC and Rae1 are RNase Mini-III required for the maturation of 23S rRNA and ribosome-associated A site endoribonuclease, respectively (61,62). As our suppressor screen identified additional mutations related to transcription, we assumed that these translation-specific RNases might not be relevant for the suppression of the *rny* deletion. Therefore, we hypothesized that the duplication of the genes encoding the main three subunits of RNA polymerase was the major reason for suppression, whereas the additional point mutations in *greA, rpoE*, or *cspD* might provide just a condition specific advantage in conjunction with the duplication of the *rpoA, rpoB*, and *rpoC* genes (see Fig. 1B).

### Genomic separation of the genes encoding the core subunits of RNA polymerase

To test the idea that simultaneous duplication of all three genes for the RNA polymerase core subunits is the key for the suppression of the loss of RNase Y, we decided to interfere with this possibility. The duplicated region is located between two highly conserved *rrn* gene clusters which may facilitate the duplication event (see Fig. 2). Therefore, we attempted to separate the core RNA polymerase genes by relocating the *rpoA* gene out of this genomic region flanked by the *rrn* operons. We assumed that if RNA polymerase was indeed the key to the original suppression, such a duplication would not be likely in the new background with relocated *rpoA*, since simultaneous duplication of all three RNA polymerase subunit genes would be disabled there. For this purpose, the *rpoA* gene kept under the control of its natural promoter P*rpsJ* was placed between the *dgk* and *yaaH* genes, and the original copy of *rpoA* was deleted (see Fig. 2, Materials and Methods for details). We then compared the growth of the wild type strain 168 and the strain with the relocated *rpoA* GP2903 using a drop-dilution assay. No differences were observed, thus excluding a possible negative impact of the *rpoA* relocation on *B. subtilis* physiology (see Supplemental Fig. S2).

**Figure 2.**
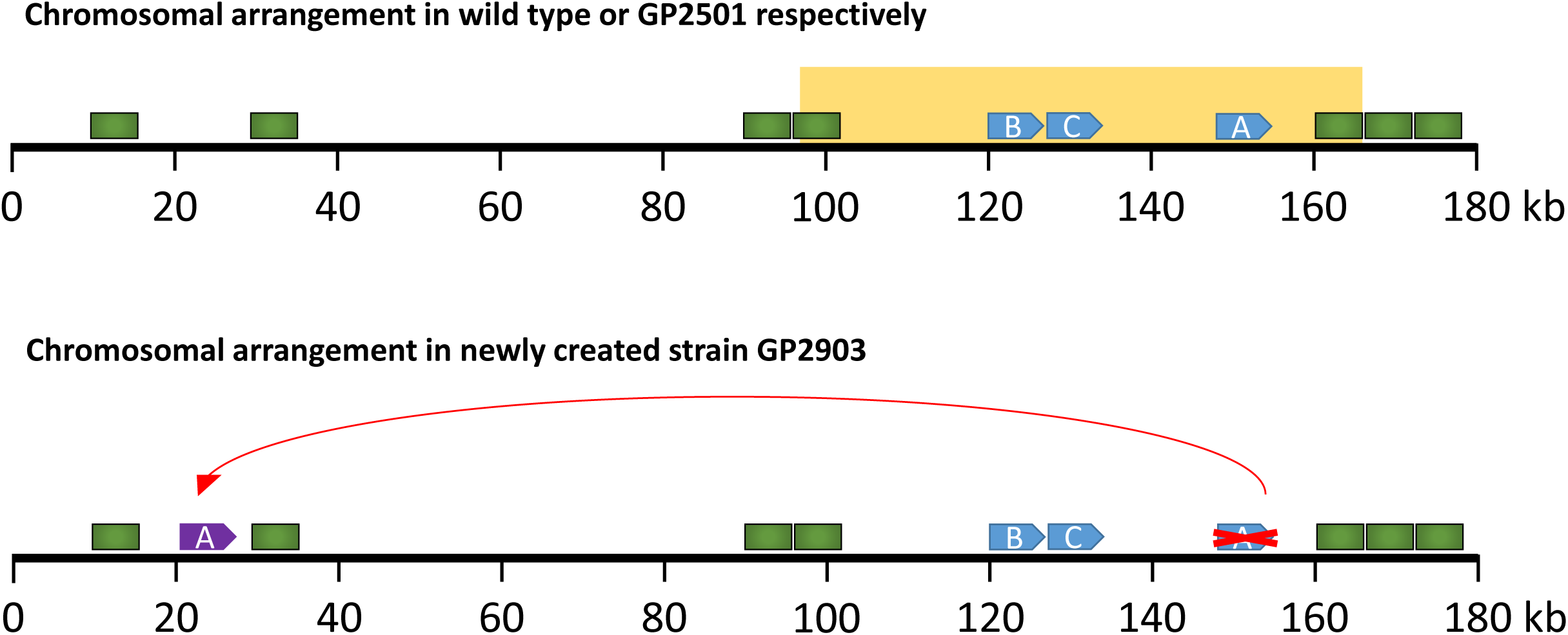
Genomic separation of the *rpoA* and *rpoBC* genes. Schematic representation of the first 180 kb of the *B. subtilis* chromosome. rRNA operons are depicted as green rectangles, RNA polymerase genes *rpoA, rpoB, rpoC* as blue arrows, and the relocated *rpoA* as a purple arrow. The orange box indicates the region which was duplicated in the suppressors of *rny* strain GP2501.

Strain GP2903 was then used to delete the *rny* gene, and to isolate suppressor mutants. Indeed, even with the genomically separated RNA polymerase genes, suppressor mutations appeared upon the deletion of the *rny* gene encoding RNase Y. There were three possibilities for the outcome of the experiment. First, the same genomic region as in the original suppressors might duplicate thus falsifying our hypothesis. Second, both regions containing the *rpoA* and *rpoBC* genes might be duplicated. Third, in the new genetic background completely new suppressing mutations might evolve. Two of these suppressor mutants were subjected to whole genome sequencing. None of them had the duplication of the *ctsR-pdaB* region as in the original suppressors. Similarly, none of the mutants had the two regions containing the *rpoA* and the *rpoBC* genes duplicated. Instead, both mutants had point mutations in the RNA polymerase subunit genes that resulted in amino acid substitutions (GP2912: RpoC, R88H; GP2913: RpoB, G1054C; see Table 1). A mutation affecting RNA polymerase was also evolved in one strain (GP2915) not subjected to whole genome sequencing. In this case, the mutation resulted in an amino acid substitution (G45D) in RpoC.

An analysis of the localization of the amino acid substitutions in RpoB and RpoC revealed that they all affect highly conserved amino acid residues (see Fig. 3A). G1054 of RpoB and G45 of RpoC are universally conserved in RNA polymerases in all domains of life, and R88 of RpoC is conserved in the bacterial proteins. This high conservation underlines the importance of these residues for RNA polymerase function. The mutations G45D and R88H in RpoC affect the N-terminal β’ zipper and the zinc-finger like motif of the β′ subunit, respectively, that are required for the processivity of the elongating RNA polymerase (63,64). G1054C in RpoB is located in the C-terminal domain of the β subunit that is involved in transcription termination (65). In the three-dimensional structure of RNA polymerase, these regions of the β and β′ subunits are located in close vicinity opposite to each other in the region of the RNA exit channel which guides newly transcribed RNA out of the enzyme (see Fig. 3B, 64), and they are both in direct contact with DNA (63).

**Figure 3.**
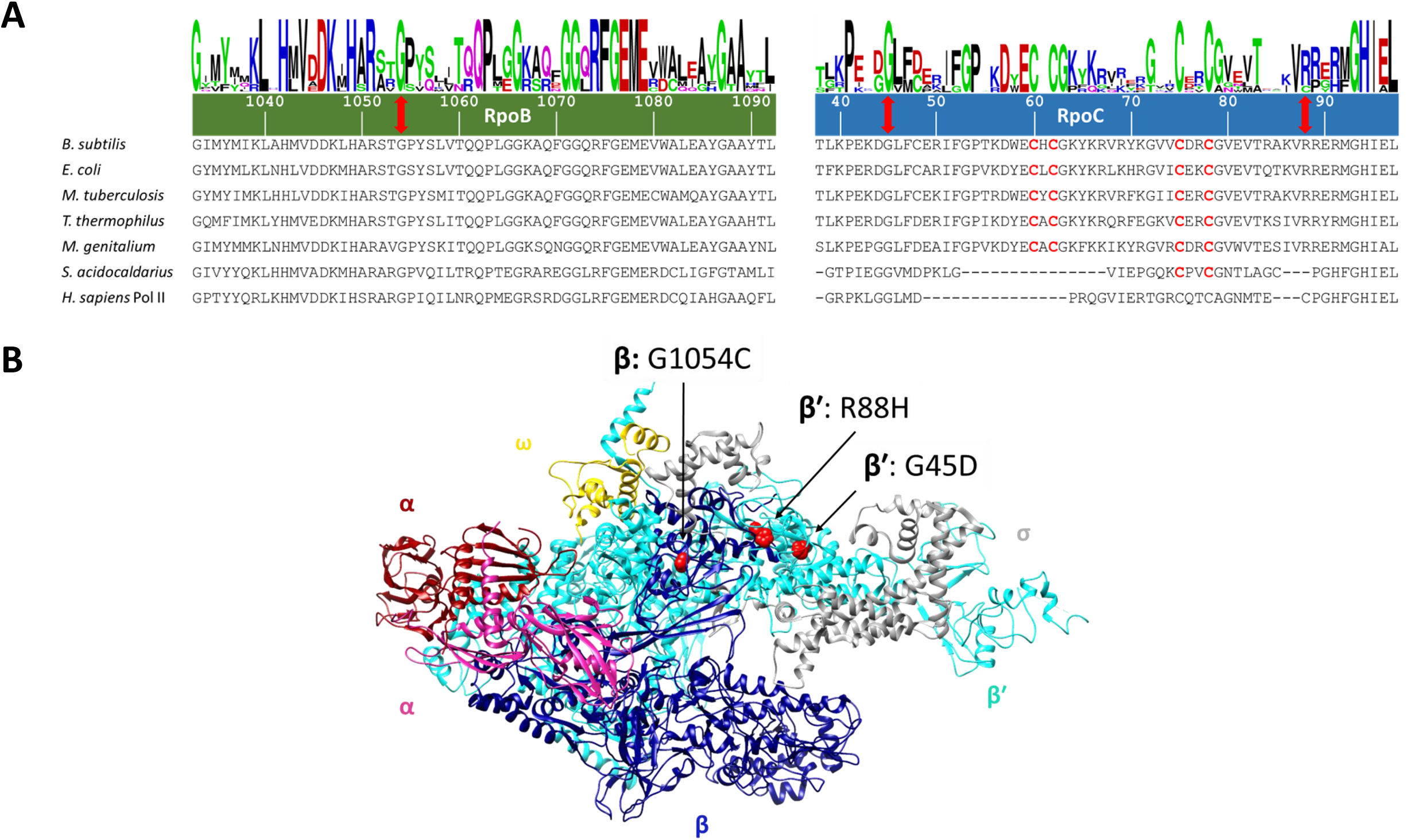
Suppressor mutations in RNA polymerase localize to evolutionary conserved regions. (A) Multiple sequence alignment of RpoB and RpoC sequences from various species, the numbering of amino acid residues is based on the *B. subtilis* sequence. The positions of mutations are indicated with red double head arrows, conserved cysteines involved in Zn-finger formation are shown in red. Logos were created as described (90). Abbreviations: *B. subtilis, Bacillus subtilis*; *E. coli, Escherichia coli*; *M. tuberculosis, Mycobacterium tuberculosis*; *T. thermophilus, Thermus thermophilus*; *M. genitalium, Mycoplasma genitalium*; *S. acidocaldarius, Sulfolobus acidocaldarius*; *H. sapiens, Homo sapiens*. (B) Localization of the mutations (indicated as red spheres) in the RNA polymerase shown at their corresponding position in the structure of *T. thermophilus* (PDB ID: 1IW7; 91). The two α subunits are shown in dark red and violet, respectively, the ß subunit is shown in dark blue, ß’ in cyan, ω in gold and the s subunit is shown in grey. The image was created using UCSF Chimera (92).

The fact that several independent mutations affecting RNA polymerase were obtained in the suppressor screen strongly supports the idea that RNA polymerase is the key for the suppression. As the mutations affect highly conserved residues, they are likely to compromise the enzyme’s activity. Based on the structural information, the mutations might weaken RNA polymerase-nucleic acid interactions and therefore, destabilize the transcription elongation complex which may result in increased premature termination and reduced RNA polymerase processivity. However, RNA polymerase is essential, therefore the mutations cannot inactivate the protein completely.

### A pre-existing duplication of the genomic region containing *rpoA* and *rpoBC* is fixed in response to the deletion of *rny*

The screen for suppressor mutations that facilitate growth of strains lacking RNase Y yielded two classes of mutants: the first set harboured mutations in genes involved in transcription (*greA, rpoE*, or *cspD*) in addition to a duplication of the chromosomal region encoding the core subunits of RNA polymerase. The second class had point mutations affecting the β or β’ subunits of RNA polymerase that result in strongly decreased transcription activity. At a first glance, these results seem to be conflicting. Considering RNA degradation as the function of RNase Y, it seemed plausible that the selective pressure caused by deletion of *rny* should result in alleviating the stress from mRNA accumulation. This seems to be the case in the second class of suppressors (see below), whereas the logic behind the duplication seems to be less obvious. Importantly, we noticed that this duplication was always accompanied by one of the other aforementioned mutations affecting transcription. In an attempt to determine the order of the evolutionary events in these suppressors we established a method to detect the presence of the duplication without whole genome sequencing. For this, we made use of a pair of oligonucleotides that binds to the *pdaB* and *ctsR* genes giving a product of about 10 kb, if the region is duplicated or amplified but no product in the absence of duplication or amplification (see Fig. 4A). This PCR product was very prominent for the strain GP2636 that is known to carry the duplication. However a band was also observed in the wild type strain 168, indicating that the duplication is present in a part of the population independent from the selective pressure exerted by the *rny* deletion (Fig. 4B).

**Figure 4.**
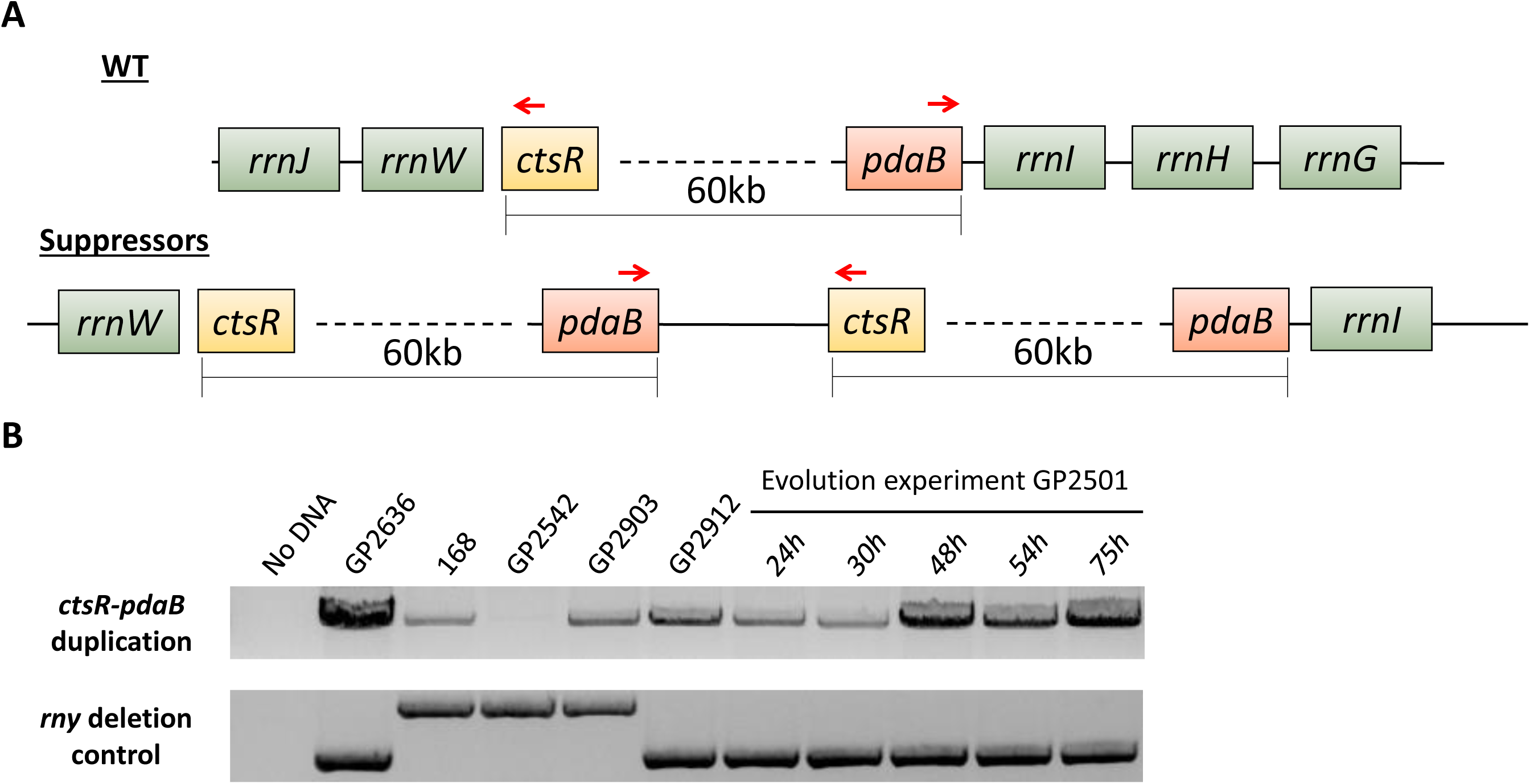
Duplication of the *ctsR-pdaB* region in suppressors of the *rny* mutant GP2501. (A) Schematic representation of the *ctsR-pdaB* region and its duplication in suppressors of GP2501. The binding sites of the oligonucleotides used for the PCR detection of the duplication is indicated by red arrows. (B) Upper panel: The PCR product obtained by PCR using primers binding to *pdaB* and *cts*R genes indicating presence of the duplication. Lower panel: The PCR product for the amplification of the *rny* region. Note the 5 µl of the PCR product were loaded in the upper panel, and 1 µl in the lower panel.

It is well-established that genomic duplications or amplifications occur frequently in bacterial populations, even in the absence of selective pressure (66). In *Salmonella typhimurium, rrn* operons have been shown to be a hotspot of gene duplications or amplifications (67). Since evolution of such a genomic duplication is dependent on homologous recombination, we performed the PCR also on the *recA* mutant GP2542, which is defective in homologous recombination and thus unable to amplify chromosomal regions (44,45). Indeed, in this case we did not obtain even a faint band. Interestingly, the genomic duplication can also be observed in cells having the core subunits of RNA polymerase at distinct genomic regions (GP2903). For the derived suppressor mutant GP2912 that carries a point mutation in *rpoC*, the band indicating the presence of the duplication was also detectable by PCR analysis although the duplication could not be detected by genome sequencing. This apparent discrepancy is most easily resolved by assuming that the duplication was present only in a small subpopulation (as observed for the wild type strain) and therefore only detectable by the very sensitive PCR assay.

Obviously, the different genomic and genetic backgrounds of the *rny* mutants generate distinct selective forces: While the duplication is not fixed in strains with separated *rpo* genes, it seems to become fixed in the suppressor mutants that have the *rpo* genes in one genomic region. To investigate the order of evolutionary events, we cultivated the *rny* mutant strain GP2501 for 75 hours and monitored the status of the *rpoA-rpoBC* chromosomal region by PCR (see Fig. 4B). The initial sample for the *rny* mutant GP2501 that was used for the experiment, already revealed the presence of the duplication in a small sub-population. This supports the finding that the duplication is present irrespective of any selection. The band corresponding to the duplicated *pdaB-ctsR* region became more and more prominent in the course of the experiment, after 75 hours it was comparable to the signal obtained with strain GP2636 that carries the duplication. As a control, we also amplified the genomic region of the *rny* gene. In the wild type strain, this PCR product has a size of 2.5 kb, whereas the replacement of *rny* by a spectinomycin resistance gene resulted in a product of 2 kb. Importantly, the intensity of this PCR product did not change during the course of the evolution experiment, thus confirming that the increased intensity of the product for the *pdaB-ctsR* region represents the spread of the duplication in the bacterial population. To verify the duplication and to check for the presence of accompanying mutations, we subjected the strain obtained in this evolution experiment after 75 hours (GP3211) to whole genome sequencing. The sequencing confirmed presence of the duplication, but did not reveal any additional suppressor mutation. Based on this result, we can assume that upon deletion of *rny* the bacteria first fixed the duplication of the *pdaB-ctsR* region and then, later, may acquire the point mutations affecting *greA, rpoE*, or *cspD*.

### Establishing the *rpoB* and *rpoC* mutations in wild type background

Based on the essentiality of transcription, we expected that the mutations in *rpoB* and *rpoC* that we have identified in the suppressor screen with the *rny* mutant and genomically separated RNA polymerase genes might adjust some of the properties of RNA polymerase. To study the consequences of these mutations for the RNA polymerase and hence also for the physiology of *B. subtilis*, we decided to introduce one of them (RpoC-R88H) into the wild type background of *B. subtilis* 168. For this purpose, the CRISPR/Cas9 system designed for use in *B. subtilis* was employed (49). As a control, we used the same procedure to introduce a mutation in the *rae1* gene, which is located nearby on the chromosome. Although this system readily allowed the introduction of a frameshift mutation (introduction of an extra T after 32 bp) in *rae1* (strain GP2901), we failed to isolate genome-edited clones expressing the RpoC-R88H variant in multiple attempts. This failure to construct the RpoC-R88H variant in the wild type background suggests that the properties of the protein are altered in a way that is incompatible with the presence of an intact RNA degradation machine.

### Mutated RNA polymerases have highly decreased activity *in vitro*

Since our attempts to study the effect of the mutations *in vivo* failed, we decided to test the properties of the mutant RNA polymerases using *in vitro* transcription. *B. subtilis* RNA polymerase is usually purified from a strain expressing His-tagged RpoC (52). However, the loss of competence of the *rny* mutant and the lethality of the *rpoC* mutation in the wild type background prevented the construction of a corresponding strain. To solve this problem, we used an approach to purify *B. subtilis* RNA polymerase from *E. coli* that had been successful before for RNA polymerase of *Mycobacterium smegmatis* (50). Briefly, plasmid pBSURNAP containing genes *rpoA, rpoB, rpoC, rpoE, rpoY*, and *rpoZ* for the RNA polymerase subunits under control of an IPTG inducible promoter was constructed in a way that each individual gene for a subunit could be cleaved out using unique restriction sites and replaced with its mutant counterpart, yielding pGP2181 (RpoC-R88H) and pGP2182 (RpoB-G1054C) (for details of the construction, see Materials and Methods). The variant RNA polymerases were expressed in *E. coli* BL21 and purified via affinity chromatography and subsequent size exclusion chromatography.

We purified the wild type and two mutant RNA polymerases (RpoC-R88H and RpoB-G1054C) and assessed their activity by *in vitro* transcription on three different templates, containing well-studied promoters of the *veg* and *ilvB* genes and the P1 promoter *of the rrnB* operon (68,69). In agreement with previous results with wild type RNA polymerase (70), this enzyme performed well on all three substrates. In contrast, the mutated variants of RNA polymerase exhibited a drastic decrease of transcription activity on all three promoters; for the RpoB-G1054C variant the transcripts were only barely detectable (Fig. 5A).

**Figure 5.**
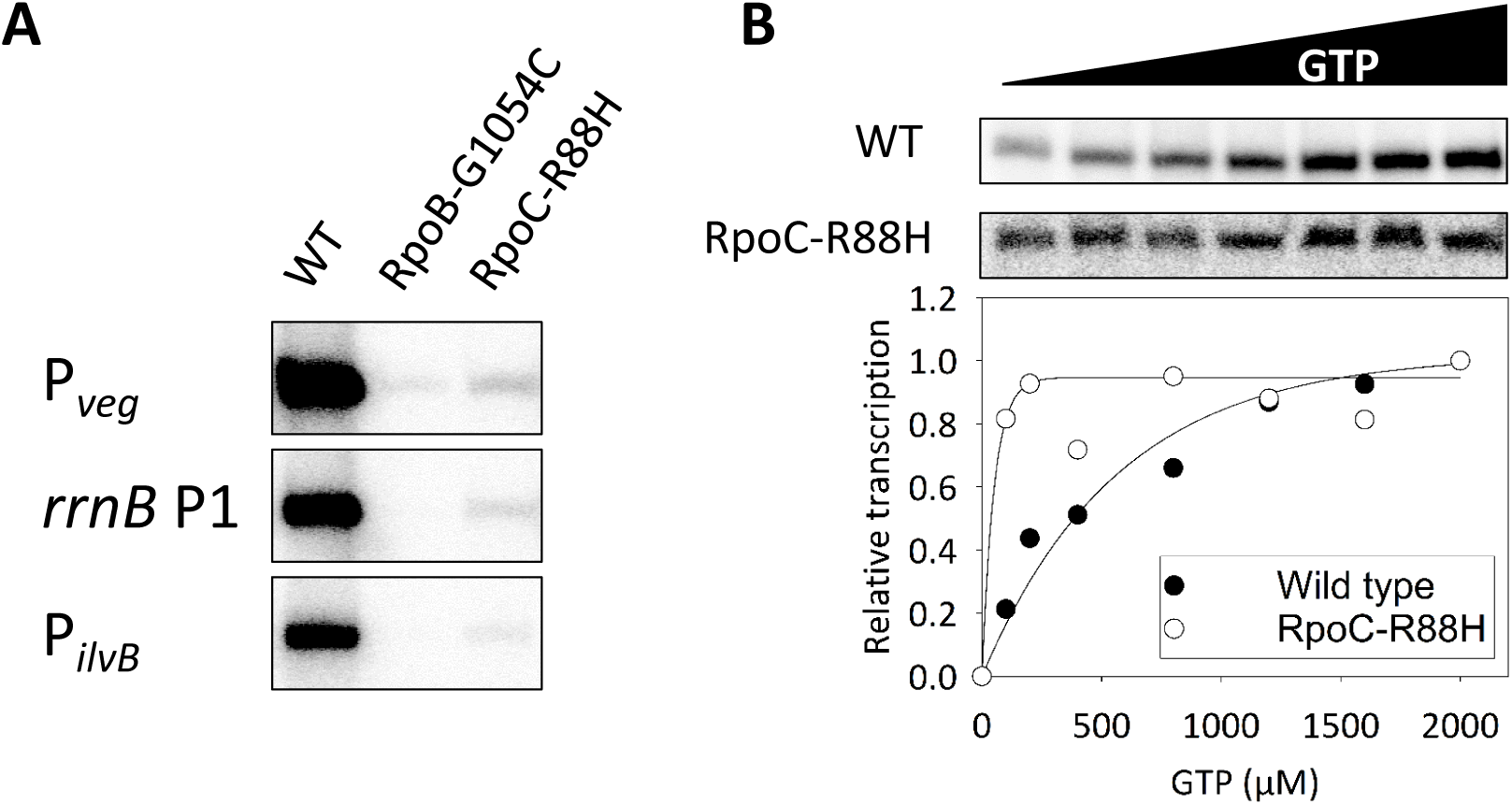
Comparison of transcriptional activity between RNA polymerase variants. (A) The RNA polymerase variants (64 nM) were reconstituted with saturating concentrations of σ^A^, e.g. in a ratio 1:10. Holoenzymes were used to initiate transcription on three different promoters as indicated. A representative image from three independent experiments is shown. (B) Primary data show transcription from the *rrnB* P1 promoter in dependence on increasing concentration of iNTP (GTP). The intensity of the transcripts generated by RNA polymerase containing RpoC-R88H was adjusted for better visibility. The relative activity of this mutant RNA polymerase was 2.5% of the activity of wild type RNA polymerase at 2,000 μM GTP. The graph shows average of two replicates normalized for maximal transcription of each polymerase (set as 1).

On many promoters, including the P1 promoter of the *rrnB* operon, *B. subtilis* RNA polymerase is sensitive to the concentration of the first transcribed nucleotide both *in vitro* and *in vivo* (68). This prompted us to compare the response of the wild type and the RpoC-R88H variant RNA polymerases to different concentrations of GTP, the initiation NTP for the *rrnB* P1 transcript. As described before, transcription with the wild type enzyme increased gradually in response to the GTP concentration (68). In contrast, the mutated variant was saturated with a relatively low GTP concentration, suggesting that this important regulatory mechanism is not functional here (see Fig. 5B).

Taken together, our results suggest that a massive reprogramming of the properties of RNA polymerase as indicated by a substantial reduction in RNA polymerase activity and its altered ability to be regulated by iNTPs allows the suppressor mutants to overcome the loss of RNase Y.

## DISCUSSION

RNases E and Y are the main players in RNA degradation in *E. coli* and *B. subtilis*, respectively. Recently, it has been estimated that about 86% of all bacteria contain either RNase E or RNase Y (or, sometimes, both) supporting the broad relevance of these two enzymes (13). While RNase E of *E. coli* is essential, conflicting results concerning the essentiality of RNase Y have been published (18,28,29,37,39). In this study, we have examined the properties of *B. subtilis* mutants lacking RNase Y due to deletion of the corresponding *rny* gene. We observed that the *rny* mutant grew poorly, and rapidly acquired secondary mutations that suppressed, at least partially, the growth defect caused by the deletion of the *rny* gene. Thus, we conclude that RNase Y is in fact quasi-essential (31) for *B. subtilis*, since the mutant cannot be stably propagated on complex medium without acquiring suppressor mutations.

A lot of effort has been devoted to the understanding of the reason(s) of the (quasi)-essentiality of RNases E and Y for *E. coli* and *B. subtilis*, respectively. Initially, it was assumed that the essentiality is caused by the involvement of these RNases in one or more key essential processing event(s) that may affect the mRNAs of essential genes as has been found for *B. subtilis* RNase III and *E. coli* RNase P (33,34,35,72,73). However, such a target was never identified. Instead, different conclusions were drawn from suppressor studies with *E. coli rne* mutants lacking RNase E: some studies reported suppression by the inactivation or overexpression of distinct genes, such as *deaD* encoding a DEAD-box RNA helicase and *ppsA* encoding phosphoenolpyruvate synthetase, respectively (74,75). In addition, the processing and degradation of the essential stable RNAs, such as tRNAs and rRNAs was shown to be an essential function of RNase E (76). Yet another study suggested that mRNA turnover is the growth-limiting factor of the *E. coli rne* mutant (71). The results presented here lend strong support to the idea that the main task of RNase Y in *B. subtilis* is the control of intracellular mRNA concentration via the initiation of mRNA degradation. Irrespective of the conditions used in the different suppressor screens, we identified a coherent set of mutations that resulted in improved growth of the *B. subtilis rny* mutant. The initial mutants carry a duplication of the chromosomal region that contains the genes for the core subunits of RNA polymerase (RpoA, RpoB, RpoC) and point mutations in *greA, rpoE*, and *cspD* that all affect transcription. If this duplication was prevented by genomically separating the RNA polymerase genes, we found suppressor mutants affecting the core subunits of RNA polymerase which result in strongly compromised transcription activity. Taken together, these findings suggest that the (quasi)-essentiality of RNases E and Y is related to their general function in initiating mRNA turnover rather than to the processing of specific RNA species. This idea is further supported by two lines of evidence: First, mutations that mimic a stringent response and therefore reduce RNA polymerase activity suppressed the growth defect of an *rne* mutant, and second, artificial expression of RNase Y or of the ribonucleases RNase J1 or J2 from *B. subtilis* partially suppressed the *E. coli* strain lacking RNase E, but only under specific growth conditions (77,78).

With the initiation of global mRNA degradation as the (quasi)-essential function of RNases E and Y in *E. coli* and *B. subtilis*, respectively, one might expect that the overexpression of other RNases might compensate for their loss. By analogy, such a compensation has been observed for the essential DNA topoisomerase I of *B. subtilis*, which could be replaced by overexpression of topoisomerase IV (44). However, in all the six suppressor mutants analysed by whole genome sequencing (Table 1), we never observed a mutation affecting any of the known RNases of *B. subtilis*. Similarly, no such compensatory mutations resulting from overexpression of other cognate RNases have been found in suppressor screens for *E. coli* RNase E. While RNase Y does not have a paralog in *B. subtilis, E. coli* possesses the two related RNases E and G. However, not even the overexpression of RNase G allowed growth of an *E. coli rne* mutant (79,80) suggesting that RNase G has a much more narrow function than RNase E and that none of the other RNases in either bacterium is capable of initiating global mRNA degradation. Interestingly, as mentioned above, RNase J1 could partially replace RNase E in *E. coli* (78), whereas it is not able to replace RNase Y in *B. subtilis*. This difference could be due to the fact that RNase J1 provides an additional pathway to initiate mRNA degradation in *B. subtilis*, which is not naturally present in *E. coli*. This idea is further supported by the observation that a *B. subtilis* strain lacking both RNases Y and J1 could never be constructed (37).

An interesting result of this study was the apparent contradiction between the isolation of suppressor mutants with increased copy number of core RNA polymerase subunit genes in one setup, and the isolation of mutants that exhibited severely reduced RNA polymerase activity in the other setup. One might expect that the increased copy number of RNA polymerase core subunit genes would result even in increased transcription. However, the outcome may just be the opposite: The RNA polymerase is a complex multi-protein machine that contains several important proteins in addition to the core subunits. As these factors, including the sigma factor and other subunits like RpoE, RpoY and RpoZ (54,58,81,82,83) as well as transcription factors like GreA and NusA (56,84) bind to the RNA polymerase via the core subunits, the perturbation of the normal evolved equilibrium between the RNA polymerase core subunits and transcription factors is likely to result in the formation of abortive incomplete complexes that are not fully active in transcription. To obtain a quantitative estimate for the formation of incomplete complexes, we estimated the stoichiometry of the complexes in the wild type from proteomics data (27). These data indicate that GreA and the RpoZ subunit are present in excess of core RNA polymerase, but not NusA, σ^A^ as well as the RpoE and RpoY subunits (Fig. 6A). Assuming that Sigma and NusA bind to the core RNA polymerase subsequently in different stages of transcription and that all other components bind independently, we estimate that 90% of all RNA polymerases are complete with GreA, RpoZ and either σ^A^ or NusA. This fraction is strongly reduced (to 23%) if the core subunits are duplicated. Instead a variety of incomplete complexes is expected (Fig. 6B). Complexes that also contain a RpoE and RpoY subunit are reduced even more strongly, from 59% to 8%. Thus, a duplication of the core subunit genes is indeed expected to result in a strong decrease of the transcription activity.

**Figure 6.**
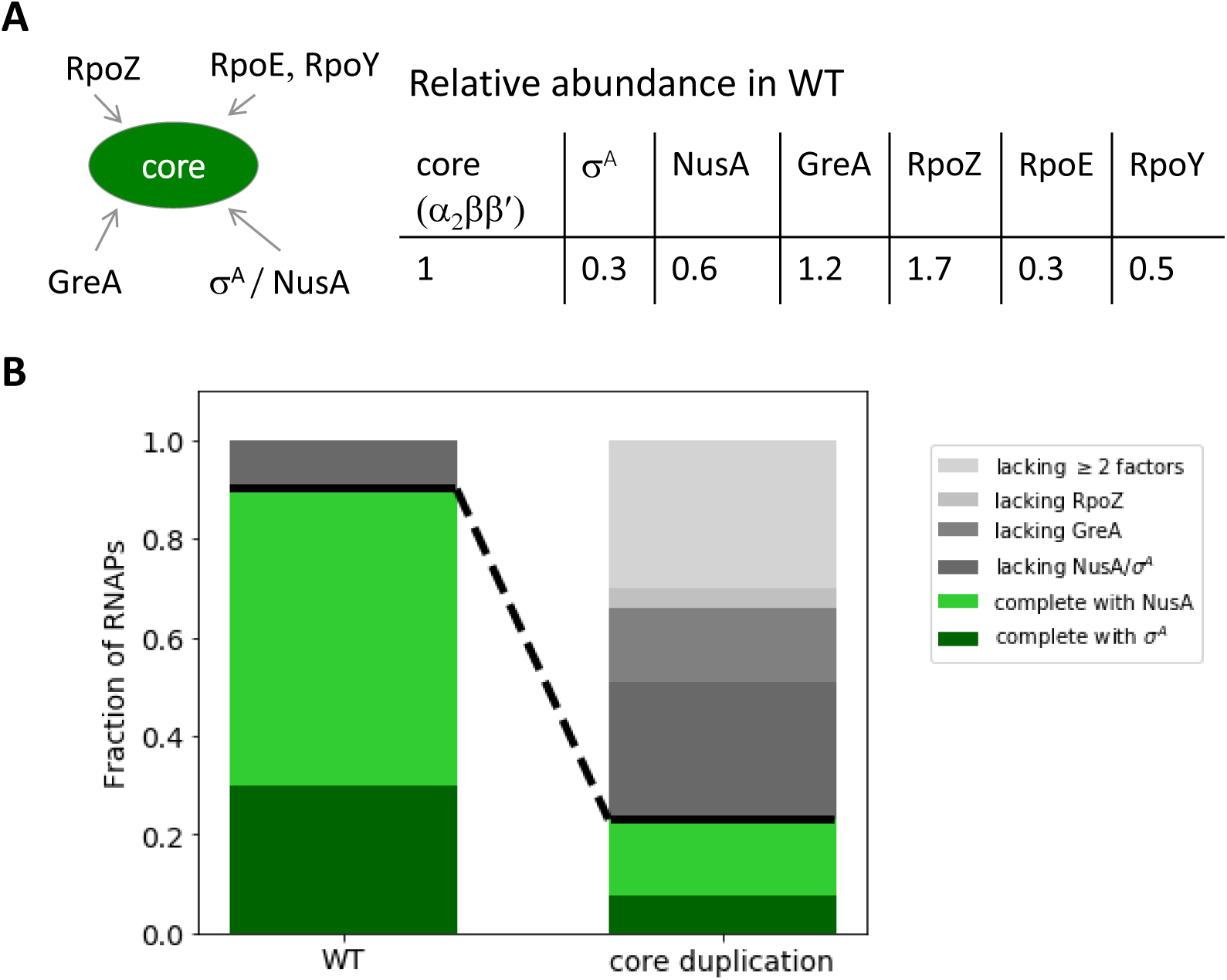
The duplication of the genes for core RNA polymerase is likely to result in the formation of incomplete RNA polymerase complexes. (A) Relative abundance/stoichiometry of RNA polymerase subunits and associated factors from proteomics data (27). (B) Fractions of core RNA polymerase in different complete (green) and incomplete (grey) complexes estimated based on the relative abundance in (A) for the wild type and for the core duplication strain, where the relative abundance of core subunits is doubled compared to all other subunits.

In each organism, an optimal trade-off between RNA synthesis and degradation must be adjusted to allow optimal growth. Obviously, the loss of the major RNA decay-initiating enzyme will bring this adjustment out of equilibrium. This idea is supported by the observation that reduced RNA degradation in *B. subtilis* is accompanied by the acquisition of mutations that strongly reduce transcription activity of the RNA polymerase. Actually, the reduction of activity was so strong that it was not tolerated in a wild type strain with normal RNA degradation. This indicates that the suppressor mutants have reached a new stable equilibrium between RNA synthesis and degradation, which, however, is not optimal as judged from the reduced growth rates of the suppressor mutants as compared to the wild type strain. It has already been noticed that generation times and RNA stability are directly related (9,85). This implies that a stable genetic system requires a balance between transcription and RNA degradation to achieve a specific growth rate. In bacteria, rapid growth requires high transcription rates accompanied by rapid RNA degradation. The association between RNA polymerase and components of the RNA degrading machinery, as shown for *B. subtilis* and *Mycobacterium tuberculosis* might be a factor to achieve this coupling between RNA synthesis and degradation (83,86).

In conclusion, our study has revealed that the initiation of mRNA degradation to keep the equilibrium between RNA synthesis and degradation is the function of RNase Y that makes it quasi-essential for *B. subtilis*. In addition to RNase Y, RNase J1 is also quasi-essential for this bacterium. In the future, it will be interesting to understand the reasons behind the critical role of this enzyme as well in order to get a more comprehensive picture of the physiology of RNA metabolism.

## Supporting information

Table S1 Oligonucleotides

## ACKNOWLEDGEMENTS

We are grateful to Alžbeta Rabatinová, Victoria Keidel, Janek Meißner and Fabian Commichau for providing strains and to Gabriele Beyer, Melanie Heinemann and Sarah Teresa Schüßler for technical support. We thank Jonas Jennrich for helpful discussions.

## AUTHOR CONTRIBUTIONS

Design of the study: MB, KG, LK and JS Experimental work: MB, SW, KG, DK, SK and HŠ Sequence analysis: MB, AP, and RD

Wrote the paper: MB, LK, and JS

