## Supplemental Figures for "Quasi-essentiality of RNase Y in *Bacillus subtilis* is caused by its critical role in the control of mRNA homeostasis"

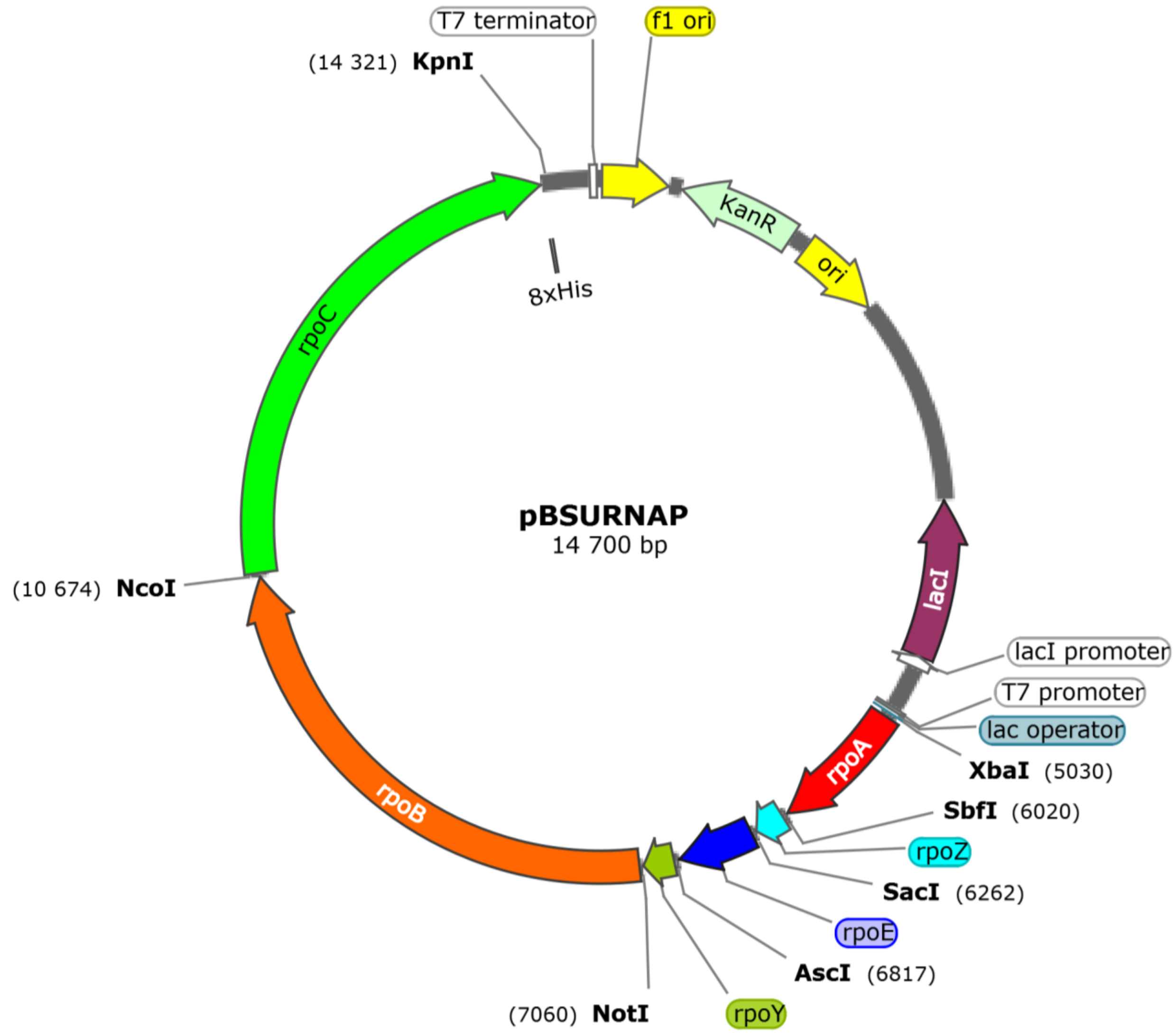


**Figure S1. Plasmid map of pBSURNAP used for the expression of the core RNA polymerase.**

Benda et al., Fig. S1

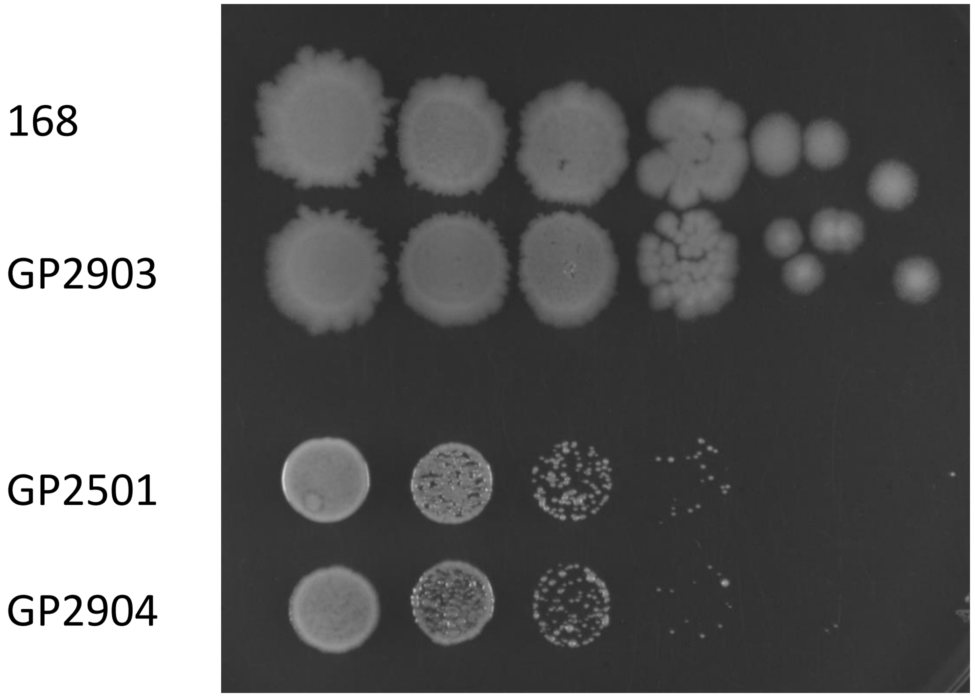


**Figure S2. Relocation of *rpoA* does not affect growth.** Serial drop dilutions comparing growth of the wild type 168, the wild type strain with relocated *rpoA* GP2903, and their respective *rny* deletion strains GP2501 and GP2904 on a LB plate at 37° C. The picture was taken after 18h of incubation.
