## Supplementary material for "Quasi-essentiality of RNase Y in *Bacillus subtilis* is caused by its critical role in the control of mRNA homeostasis": Table S1 Oligonucleotides

Table S1. Oligonucleotides used in this study

| **Primer** | **Sequence**  Restriction sites are underlined  His-tag sequences are italic  Homologous bases for joining PCR are bold |
| --- | --- |
| **Detection of *ctsR-pdaB* region duplication** | |
| MB30 | 5′-CTGTATGTCTTTGACCCCTAACTTTTC |
| MB208 | 5′-CCTCTTTCGCTTGTAAATCTGGT |
| **Control of *rny* deletion** | |
| NC9 | 5′-CTATGAAAAGATGTTTACGCCAGGG |
| ML101 | 5′‑CTGCAAATTAATGACTGCTAGTTCTT |
| **Construction of pBSURNAP and its derivatives** | |
| LK#2684 | 5′-GGTCTAGAGCGGCCGCTTTAAGAAGGAGATATATCTATGACAGGTCAACTAGTTC |
| LK#2685 | 5′-CGCGGATCCGGTACCCCATGGCGCGCAAGTTCTTTTGTTACTACATCG |
| LK#2686 | 5′-GCGCCATGGTGGCTCGGGTGCAATGCTAGATGTGAACAATTTTGAG |
| LK#2687 | 5′-GCGGTACC*TTAGTGATGGTGATGGTGATGGTGATG*TTCAACCGGGACCATATCG |
| MB169 | 5′-AAAGCGGCCGCTTTAAGAAGGAGATATATCTATGACAGGTCAACTAGTTCAGTATGGAC |
| MB170 | 5′-AAACCATGGCGCGCAAGTTCTTTTGTTACTACATCGCGTTCAA |
| MB167 | 5′-AAACCATGGTGGCTCGGGTGCAATGCTAGATGTGAACAATTTTGAGTATATGAAC |
| MB168 | 5′-AAAGGTACC*CTAGTGATGGTGATGGTGATGGTGATG*TTCAACCGGGACCATATCGT |
| **Deletion of *cspD* gene** | |
| MB17 | 5′-CGCCGAACTGGAAGAGTCATTCC |
| MB18 | 5′-**CCTATCACCTCAAATGGTTCGCTG**GTTGAACCATTTTACTTTACCGTTTTGCAT |
| MB19 | 5′-**CCGAGCGCCTACGAGGAATTTGTATCG**GTAATCGTGGACCTCAAGCTTCTAATGTTG |
| MB20 | 5′-GAAGCACTCCTTGAATCGCTGAAGC |
| kan-fwd | 5′-CAGCGAACCATTTGAGGTGATAGG |
| kan-rev | 5′-CGATACAAATTCCTCGTAGGCGCTCGG |
| MB21 | 5′-GGCGAACTTGTCGATGAACATCAG |
| MB22 | 5′-GGCAGCTGGCCTTGTTATGATC |
| **Deletion of *rny* gene** | |
| ML47 | 5′-GAAGAATCTGCTTACACATACATCG |
| KG409 | 5′-**GACTGTGTTTTATATTTTTCTCGTTCAT**ACTTTCACCTCCTCTTGCTATGAACT |
| KG410 | 5′-**CCGAGCGCCTACGAGGAATTTGTATCG**AGTGATGCGCTAAGCATCACTTTATTTTTTTG |
| NP60 | 5′-GCAGACACATACTCTCCCACTTTTACACTGCTGACAT |
| KG411 | 5′-**ATGAACGAGAAAAATATAAAACACAGTC** |
| CZ68 | 5′-**CGATACAAATTCCTCGTAGGCGCTCGG**TTACTTATTAAATAATTTATAGCTATTG |
| KG414 | 5′-GTCGGTTCATCACAAAAAGCGCTGAT |
| NP61 | 5′-AGTATTGGTACACACATGAGATTTTCCTGTTAG |
| NC16 | 5′-CTGCCACTGAATTTGGACTCG |
| ML101 | 5′‑CTGCAAATTAATGACTGCTAGTTCTT |
| **Construction of the strain GP2909 with relocated *rpoA* gene** | |
| SW17 | 5′-GACATTGTCCCTTTATCAGC |
| SW18 | 5′-**GGGGTGTGAGCTGAATTC**CTGCTGTCTGATCAATTTAATG |
| SW19 | 5′-**CCGAGCGCCTACGAGGAATTTGTATCG**CCCCCATGAAAAAAAGAC |
| SW20 | 5′-CGAATCAAATGCTTATTTGG |
| SW21 | 5′-GAATTCAGCTCACACCCC |
| SW40 | 5′-**TGGTTTTTCAATCTCGAT**CATTATTTTCCCTCCTTTTC |
| SW23 | 5′-ATGATCGAGATTGAAAAACCA |
| SW24 | 5′-**CCTATCACCTCAAATGGTTCGCTG**TCAATCGTCTTTGCGAAG |
| cat-fwd (kan) | 5′-**CAGCGAACCATTTGAGGTGATAGG**CGGCAATAGTTACCCTTATTATCAAG |
| cat-rev (kan) | 5′-**CGATACAAATTCCTCGTAGGCGCTCGG**CCAGCGTGGACCGGCGAGGCTAGTTACCC |
| SW25 | 5′-GATCATAATCTTCAATGCGAAG |
| SW27 | 5′-GAACAACCACAAATGACATC |
| SW28 | 5′-GTGATCTGTGAAAATCCAAAG |
| SW29 | 5′-**CCTATCACCTCAAATGGTTCGCTG** TACTTAAAACCCTCCTTCAAAAC |
| SW30 | 5′-**ATATTTTACTGGATGAATTGTTTTAGTAA** CTAGTTTCCCTTGTGAACTAGG |
| SW31 | 5′-CAACTCTCTGCTTTTGGC |
| kan-fwd | 5′-CAGCGAACCATTTGAGGTGATAGG |
| kan-rev w/o T. | 5′-TTACTAAAACAATTCATCCAGTAAAATAT |
| SW32 | 5′-GCGATGTTCAAAGTTGAAC |
| SW33 | 5′-CATATTTTTTACCGCCATTCA |
| SW41 | 5′-TTGTCAAGTGAAGGCGCGCTAT |
| mls-rev (kan) | 5′-**CGATACAAATTCCTCGTAGGCGCTCGG**GCCGACTGCGCAAAAGACATAATCG |
| SW42 | 5′-AGTGAAGGAAAAGGGATG |
| SW43 | 5′-**CCGAGCGCCTACGAGGAATTTGTATCG**GTTCTGTAGATTCACTCCGA |
| SW44 | 5′-**ATAGCGCGCCTTCACTTGACAA**GAGACTTAGATTAAGTTGACGC |
| SW45 | 5′-ACTGTCAATATAGCATAAATTCC |
| **Construction of plasmid pGP2542** | |
| KG412 | 5′-AAAGGATCCATGGCACAAGAGAAAGTTTTTCCTATG |
| KG413 | 5′-TTTGTCGACTGAAATTTTCACAATTTTCACGAGCATTTC |
| **Construction of plasmid pLK502** | |
| LK#125 | 5′-GGGAATTCATGGATTGCAAGATGATCTG |
| LK#127 | 5′-CCAAGCTTAGACCGAACTCATATTACGCCGC |
| **Construction of pGP2825 for CRISPR editing ^1^** | |
| SW69 | 5′-AAGGCCAACGAGGCCCTTACTCACTTGTTAC |
| SW70 | 5′-AAGGCCTTATTGGCCTCTTGAAGCATACG |
| SW73 | 5′-aAGgATGGGaCACATTGAACTGGCTG |
| SW74 | 5′-tCCCATcCTtTCACGAtGGACTTTAGC |
| SW93 | 5′-(p)TACGTAAAGTCCGTCGTGAGAGAA |
| SW94 | 5′-(p)AAACTTCTCTCACGACGGACTTTA |
| **Construction of pGP2826 for CRISPR editing ^1^** | |
| SW81 | 5′-AAGGCCAACGAGGCCCAAGCACAGCAAGTGATT |
| SW82 | 5′-aATGTTGTAgCCaTCgACcAACAGGATATCCATGGGT |
| SW83 | 5′-gGTcGAtGGcTACAACATtGATTGGAGCC |
| SW84 | 5′-AAGGCCTTATTGGCCCCTGAACAATATCCTCTCTG |
| SW85 | 5′-(p)TACGTGGATATCCTGTTAGTAGAC |
| SW86 | 5′-(p)AAACGTCTACTAACAGGATATCCA |
| **Sequencing primers** | |
| ***cspD*** |  |
| KG227 | 5´- GAAGGAATCAGAAATGATGACCGCCA |
| KG228 | 5´- CGCTGTTTCCACCGCTAGTTCCA |
| ***rpoE*** |  |
| MB5 | 5´-CACGCAAATCTATGAAGGCACTC |
| MB6 | 5´-GCTACAATACCCTTTCCAAGTGAG |
| ***adeR*** |  |
| MB206 | 5´-GTCTTGCCTCCGATGACTTTC |
| MB207 | 5´-GCGCCTGTTTCAACCAGCA |
| ***greA*** |  |
| KG384 | 5´-GAACGAGGACTGCCCTGTGTTCTC |
| KG385 | 5´-CTGCCAGCTTCATTCGTTTCGATATCTTC |
| ***rpoB*** |  |
| MB9 | 5´-GAAGGCGTATCTGAGCGTGACG |
| MB176 | 5´-GAATCGCCTCTTCAATCAGAGAC |
| MB177 | 5´-TGGATGTATCGCCTAAGCAGGTT |
| SW63 | 5´-GATTCTTCCTGAAGAGGATATG |
| KG422 | 5´-GGATCAGTTACAACGTAAGAAGC |
| ***rpoC*** |  |
| MB108 | 5´-GCGCTCAATTGTTTCAGTTCCTTC |
| MB175 | 5´-CAATTGTCCCGCAGTATAAGCTG |
| SW77 | 5´-GATACCGCTCTTAAAACTGC |
| SW87 | 5´-GAAACAAGCCTTCTTGGA |
| SW88 | 5´-CGTACCATCACGTATGAAC |
| **pJOE8999** |  |
| SW79 | 5′-GAATACCGGTTGCATCTG |
| SW80 | 5′-AGATTATTGAGCAAATCAGTG |
| **pGP2542** |  |
| M13fwd | 5′-GTAAAACGACGGCCAGTG |

^1^ Phosphorylated primers are indicated with (p). Mutation position are written in lower case.
